# Genome assembly of a Mesoamerica-derived *Phaseolus lunatus* cultivar enables the identification and comparative genomic analysis of a complex resistance locus

**DOI:** 10.1101/2020.08.19.257105

**Authors:** Randall J. Wisser, Sara J. Oppenheim, Emmalea G. Ernest, Terence T. Mhora, Colin Scanlan, Michael D. Dumas, Nancy F. Gregory, Thomas A. Evans, Nicole M. Donofrio

**Author notes:** FMC Stine Research Center, 1090 Elkton Rd, Newark, DE 19711. **Corresponding Author:** Nicole M. Donofrio, 152 Townsend Hall, 531 S. College Avenue, Department of Plant and Soil Sciences, University of Delaware, Newark, Delaware, 19716.

## Abstract

Lima bean, *Phaseolus lunatus*, is closely related to common bean and is high in fiber and protein, with a low glycemic index. Lima bean is widely grown in the state of Delaware, where late summer and early fall weather are conducive to pod production. The same weather conditions also promote diseases such as pod rot and downy mildew, the latter of which has caused previous epidemics. A better understanding of the genes underlying resistance to this and other pathogens is needed to keep this industry thriving in the region. Our current study sought to sequence, assemble and annotate a commercially available cultivar called Bridgeton, which could then serve as a reference genome, a basis of comparison to other *Phaseolus* taxa and a resource for identification of potential resistance genes. Combined efforts of sequencing, linkage and comparative mapping analysis resulted in a 625 Mb genome assembly, as well as a better understanding of a locus underlying resistance to the current downy mildew race in the field.

## INTRODUCTION

Lima bean (*Phaseolus lunatus* L.) was independently domesticated in Mesoamerica and the Andes (Motta-Aldana *et al*. 2010; Serrano-Serrano *et al*. 2011) and is now cultivated throughout the world. Lima bean is high in fiber and protein, and has a slow digestion rate of starch, making it a healthy, low glycemic index food (Bello-Perez *et al*. 2007).

In the United States, lima bean is grown predominantly in the Mid-Atlantic Region (MAR), specifically New Jersey, Delaware and Maryland. In Delaware alone, lima bean production is approximately a $6 million industry (Evans *et al*. 2007). The same conditions that are conducive to robust pod production however, are also conducive to development and proliferation of several oomycete pathogens, including downy mildew caused by *Phytophthora phaseoli* and pod rot caused by *P. capsici* (Davidson *et al*. 2002; Evans *et al*. 2007; Davidson *et al*. 2008). In 1998, a new race of *P. phaseoli* emerged to decimate lima bean crops in Delaware, prompting increased studies on this important plant, and its pathogens (Evans *et al*. 2002; Evans *et al*. 2007). In 2016, Mhora et al. used bulked segregant analysis to map the resistance locus effective against the predominant field race F of *P. phaseoli*. Collinearity analysis with the closely related and recently sequenced species *Phaseolus vulgaris* (common bean) revealed a Resistance (R) gene-dense region with 38 of the 110 genes annotated as R-gene homologs (Goodstein *et al*. 2012; Schmutz *et al*. 2014).

While the most closely linked markers to race F resistance were 1.39 Mb apart, there are many predicted genes in this region, making it difficult to know which is responsible for race F resistance. Further analysis of the genome was merited not only to improve marker proximity to the gene or genes mediating resistance to race F, but also for further exploration of the genome to reveal additional resistance gene-dense regions.

In addition to R-gene mediated disease resistance, research into lima bean diversity and the genetic basis of more complex traits, such as resistance to the pod rot pathogen and a slow mildewing phenotype observed for the cultivar Cypress (pers. comm. TA Evans), is hindered by relatively limited genomic data. In order to add to the repository of public genomic resources for lima bean, we sequenced the genome of the *P. lunatus* commercial cultivar Bridgeton, a variety of Mesoamerican origin. Further, we generated linkage and comparative maps to form larger scaffolds which aided assembly, and we performed QTL analysis to define the partial resistance associated with slow mildewing.

## MATERIALS AND METHODS

### Biological Materials

Multiple, individual seeds of Bridgeton were planted; a single plant, Bridgeton-DES4, was chosen to self-propagate seed for the reference genome. Progeny of Bridgeton-DES4 were grown in continuous darkness to generate etiolated tissue for DNA extraction. Extractions were performed using the Qiagen Maxi DNA extraction kit, followed by a DNA purification step as described by Greco *et al*. (2014) and the DNA was checked for purity using the NanoDrop ND-1000 spectrophotometer (Thermo Fisher Scientific, MA) and, quantified using both Picogreen (Thermo Fisher Scientific, MA) and QUBIT (Thermo Fisher Scientific, MA). To ensure there was no DNA fragmentation or DNA impurities, fragment analysis and agarose gel electrophoresis were used to check the quality of the high molecular weight DNA.

### Sequencing

The standardized DNA extract from a single progeny of Bridgeton-DES4 was shipped to NRGene and subjected to library construction according to their protocols. Five size-fractions were selected ranging from 470 bp to 10 kb to construct sequencing libraries following the manufacturer’s protocols (Illumina, San Diego, CA). The TruSeq DNA Sample Preparation Kit version 2 with no PCR amplification (PCR-free) was used to make replicate paired-end libraries for the 470 bp and 800 bp size fractions. The Nextera MP Sample Preparation Kit was used to make mate-pair (MP) libraries with 2-5 kb, 5-7 kb and 7-10 kb jumps. The 470 bp libraries were sequenced as 2×265 nucleotides on the Hiseq2500 v2 in rapid mode. The 800 bp libraries and part of the three MP libraries were sequenced as 2×160 bp nucleotides on the HiSeq2500 (v4 Illumina chemistry) while the remainder of the MP libraries were also sequenced as 2×150 bp on the HiSeq4000. A total of 246 Gb sequencing data (equivalent to ~360X genomic coverage, based on an estimated genome size of 0.685 Gb). All library construction and sequencing were performed at Roy J. Carver Biotechnology Center, University of Illinois at Urbana-Champaign.

### Assembly

Genome assembly was conducted using the DeNovoMAGIC™ software platform (NRGene, Ness Ziona, Israel). This is a DeBruijn graph-based assembler, designed to efficiently extract the underlying information in the raw reads to solve the complexity of the DeBruijn graph due to genome polyploidy, heterozygosity and repetitiveness. This task is accomplished using accurate-reads-based traveling in the graph that iteratively connects consecutive phased contigs over local repeats to generate long phased scaffolds (Lu *et al*. 2015; Hirsch *et al*. 2016; Avni *et al*. 2017; Zimin *et al*. 2017; Zou *et al*. 2017).

In brief, the algorithm is composed of the following steps:

1. Pre-processing: PCR duplicates, Illumina adaptor AGATCGGAAGAGC and Nextera linkers (for MP libraries) were removed. The PE 450bp 2×265 bp libraries overlapping reads were merged with minimal required overlap of 10 bp to create the stitched reads.
2. Error correction: Following pre-processing, merged PE reads were scanned to detect and filter reads with putative sequencing error (contain a sub-sequence that does not reappear several times in other reads).
3. Contigs assembly: The first step of the assembly consists of building a De Bruijn graph (kmer=127 bp) of contigs from all of the PE and MP reads. Next, PE reads were used to find reliable paths in the graph between contigs to resolve repeats and extend the contigs.
4. Scaffolds assembly: Contigs were linked into scaffolds with PE and MP information, estimating gaps between the contigs according to the expected distance of PE and MP links.
5. Fill Gaps: A final gap filling step used PE and MP links and De Bruijn graph information to detect a unique path connecting the gap edges.

### Linkage and Comparative Genomic Maps

Linkage mapping and comparative genomics approaches were used to cluster and order the assembled scaffolds into draft pseudomolecules. Linkage mapping was performed using GBS data on 163 F_2_ progeny from a cross between cultivars Cypress (slow mildewing phenotype) and Jackson Wonder. 192-plex GBS libraries were constructed following the protocol by Manching *et al*. (2017) using RASP-2.0 adapters redesigned for *Csp*6I/*Msp*I and *Csp*6I/*Taq*^α^I pairs of restriction enzymes. Samples included the F_2_ progeny along with replicate samples of the parents and F_1_ progeny as well as a negative control with no DNA. Separate libraries were constructed for *Csp6I/MspI* and *Csp*6I/*Taq*^α^I. Sequencing was performed at University of Delaware’s Sequencing and Genotyping Center on a Hiseq2500 run in rapid mode at 1×151 bp.

Processing of GBS data was performed using RedRep (https://github.com/UD-CBCB/RedRep). First, FASTQ files from two lanes of sequencing were merged for the corresponding libraries. Following quality control and barcode deconvolution, FASTQ files from the same barcode were merged and then mapped to the NRGene sequence assembly. Variants were typed using HaplotypeCaller, and the genotype matrix was filtered as follows. VCFtools (Danecek *et al*. 2011) was used to set genotype calls to missing if fewer than three reads supported the call. The resulting genotype matrix was processed in R version 3.4.1 (R Core Team 2017) with custom scripts to filter markers that: (i) had greater than 75% missing data; (ii) had inconsistent genotype calls between parental replicates; (iii) were heterozygous in either parent; (iv) did not have the expected parent-hybrid trio genotypes; and (v) had F_2_ allele frequencies less than 15% or greater than 85%.

Using sequencing scaffolds containing at least five markers, missing data were imputed using LB-Impute (Fragoso *et al*. 2016). A linkage map was constructed with QTL IciMapping software v 4.1.0.0 (Meng *et al*. 2015) (run settings: DIS [20 cM] grouping function; RECORD ordering algorithm; SARF [window size=5] rippling criterion).

Chromosomer (Tamazian *et al*. 2016) was used to align the NRgene scaffolds against the *P*. *vulgaris* reference genome (Schmutz *et al*. 2014) downloaded from JGI (“*Pvulgaris_442_v2.0.softmasked.fa.gz*”). First, while retaining softmasked sequences in the reference genome identified by analysis with RepeatMasker, LAST (Kiełbasa *et al*. 2011) was used to identify and softmask additional repeat sequences using the ‘NEAR’ seeding scheme (run settings: -uNEAR and -R11). Following guidelines for human-ape alignments (https://github.com/mcfrith/last-genome-alignments), substitution and gap frequencies were determined with “last-train” (run settings: --revsym --matsym --gapsym -E0.05 -C2) and “lastal” alignment was performed (run settings: -fMAF -K2 -m50 -E0.05 -C2, and -p corresponded to the output from last-train). Alignments between repeat sequences were discarded with “last-postmask” and a python script was used to retain only the top two LAST matches per query sequence. The resulting output was used to run the “fragmentmap” and “assembler” routines of Chromosomer. Finally, LAST was used to align the Chromosomer assembled *P*. *lunatus* genome to the *P*. *vulgaris* reference genome (run settings: -m50 -E0.05 -C2), and “last-dotplot” was used to visualize the alignment after reformatting the data accordingly.

### QTL analysis

Disease reaction to race F of downy mildew (caused by *P. phaseoli*) was determined for 384 F_2:3_ families of the Cypress X Jackson Wonder cross (163 of the F_2_ parents of these families were used to construct the linkage map; see above). Plants were grown in a greenhouse humid chamber and inoculated at emergence as described by Santamaria *et al*. (2018). All families were replicated across four separate sequential plantings (four batches across time) with five plants per family in a single pot in each round. Pots were arranged in a randomized incomplete block design with six sub-blocks augmented with repeated checks of resistant, tolerant and susceptible varieties. Individual plants were measured for lesion length on the stem and rated for the quantity of sporulation on a 1-5 scale. To account for differences in stem length, lesion length was standardized by plant height. Adjusted means for sporulation rating and height-standardized lesion length on the F_2:3_ progeny were used as an estimate of the F_2_ [1] phenotype for QTL analysis. QTL IciMapping software v 4.1.0.0 (Meng *et al*. 2015) was used to perform inclusive composite interval mapping with a LOD threshold of 2.5 and step size of 1 cM.

### Gene Annotation

Prior to structural annotation, repetitive elements were identified and masked (File S1) using RepeatMasker (Smit *et al.*) with a custom library that included all *Viridiplantae* entries from Repbase (Bao *et al*. 2015) along with repetitive elements from *P. vulgaris* (Gao *et al*. 2014). Using the masked assembly, gene predictions were generated by AUGUSTUS with the *Arabidopsis thaliana* training set (Keller *et al*. 2011; Scalzitti *et al*. 2020). The completeness of the genome assembly and AUGUSTUS gene models were analyzed with BUSCO, which measures completeness in terms of evolutionarily informed expectations of gene content (Simão *et al*. 2015). The BUSCO Embryophyta dataset (1,440 single-copy conserved genes) was used as a reference.

Functional annotation of the predicted genes was undertaken with multiple tools. First, BLASTp searches (-evalue 1e-10) were performed against a custom BLAST database that included all Viridiplantae sequences from the NCBI nr database (Wheeler *et al*. 2007; search conducted 07/12/2020). For sequences that had no hit, a second BLASTp search was performed against all sequences in the nr database. In addition, the predicted proteins for *P*. *lunatus* were compared to the *P*. *vulgaris* reference proteome (downloaded 07/12/2020 from https://www.uniprot.org/proteomes/UP000000226).

We used protein domain analysis to identify potential plant disease resistance genes (R-genes) in the genome of *P*. *lunatus*. Using NCBI’s CD-Search tool (Marchler-Bauer *et al*. 2004), we submitted the predicted *P. lunatus* protein sequences as queries to the Conserved Domain Database (search conducted online at https://www.ncbi.nlm.nih.gov/Structure/bwrpsb/bwrpsb.cgi). We filtered the CD-Search output to retain all protein that contained both NB-ARC (https://www.ebi.ac.uk/interpro/entry/InterPro/IPR002182/) and LRR (https://www.ebi.ac.uk/interpro/entry/InterPro/IPR032675/) domains, a combination that is characteristic of plant R-genes (Gao *et al*. 2018).

### Data Availability

Illumina genome sequencing data and the reference genome assembly for *P*. *lunatus* cv. Bridgeton are available at the Sequence Read Archive and NCBI database, under the BioProject accession number PRJN647124. The BioSample number is SAMN15394833. Illumina SRA entries are deposited under SRX####. The GenBank assembly accession number is GCA_####. Supplemental material available at Figshare: https://doi.org/#####.

## RESULTS AND DISCUSSION

The Mesoamerican lima bean cultivar Bridgeton was sequenced at an average depth of ~70X across the 623 Mb assembly (Table S1), approximately 91% of the 685 Mb genome estimated for *P. lunatus* (Bennett and Leitch 2012). The assembly comprised 19,316 scaffolds, 36 and 262 of which captured 50% and 85% of the expected genome size, respectively (Figure 1), with a corresponding N50 of 3.99 Mb and N85 of 0.26 M. The N50 was 5.17 Mb with respect to the sequence space (as opposed to the expected genome size). Consistent with the nucleotide composition of higher plant genomes, the GC content for lima bean was 38%, the same as its closest relative with a reference genome, common bean (Schmutz *et al*. 2014). The genome was analyzed for completeness, returning a BUSCO score of 93% (Table 1). Taken together, this study produced a near-complete assembly of lima bean for future research. However, this assembly is comprised of thousands of scaffolds which vastly exceeds the 11 (2*n* = 22) chromosomes of *P*. *lunatus* (Bonifácio et al. 2012).

**Figure 1.**
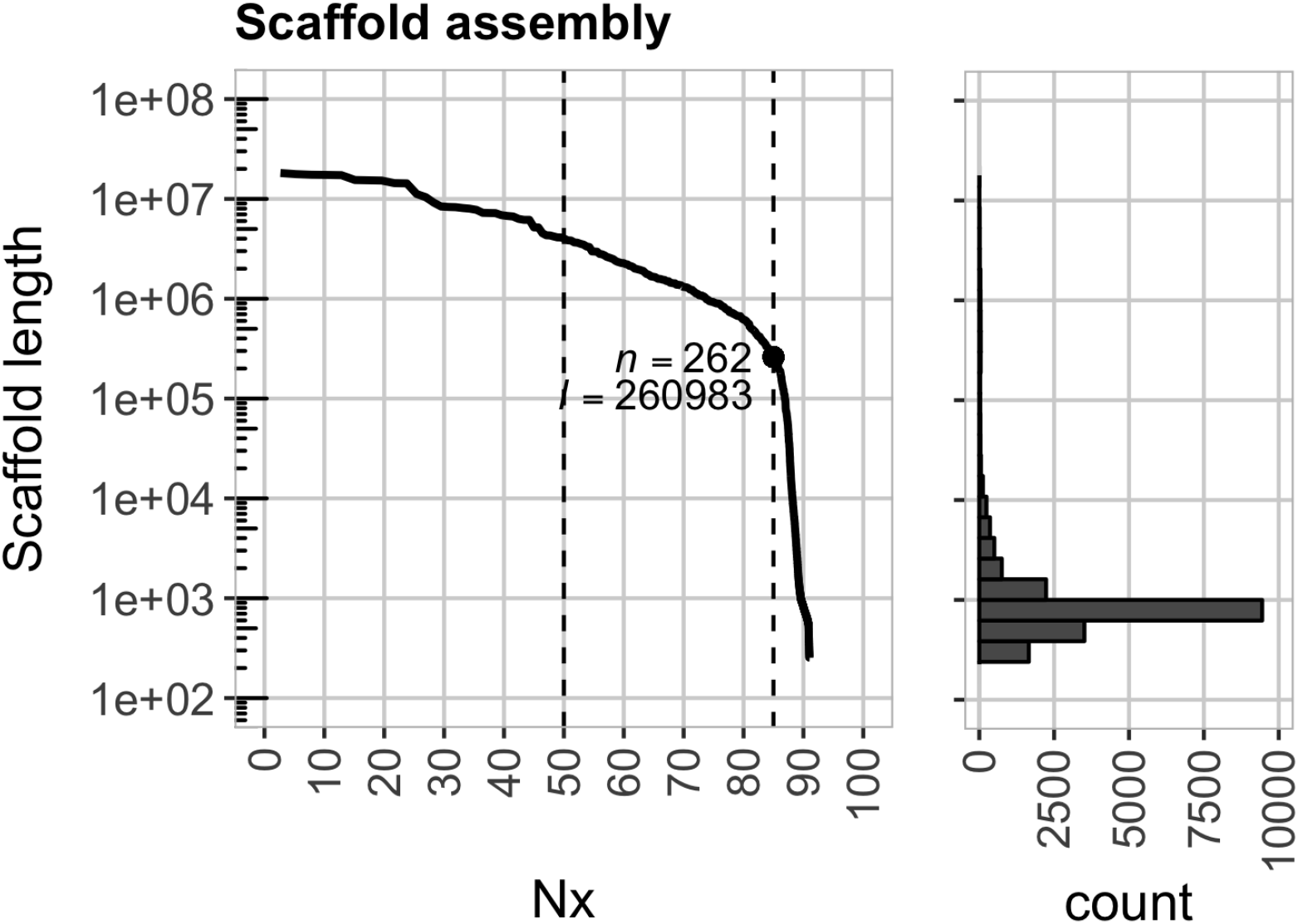
Summary statistics for the scaffold assembly of lima bean cultivar Bridgeton-DES4. (**a**) The plot shows the scaffold length as a function of the contiguity value (Nx). The N50 and N85 are marked by the dashed vertical lines where the corresponding number (*n*) and cumulative length (*l*) of scaffolds are noted. The marginal histogram plot shows the number of scaffolds at different lengths.

**Table 1.** Genome statistics and annotated gene data.

|  |  |
| --- | --- |
| Total Scaffolds | 19,316 |
| Total Contigs | 58,167 |
| Complete BUSCOs | 1341 (93.12%) |
| Scaffolds with predicted proteins | 3,417 |
| Putative Proteins | 64,541 |
| Proteins with *P. vulgaris* hit | 32,505 |
| Proteins with other Viridiplantae hit | 1,356 |
| Proteins with non-Viridiplantae hit | 2,363 |
| Proteins with no hit in nr database | 28,317 |
| Genes with predicted NB-LRR domains | 206 |

Linkage mapping and comparative genomic analysis methods were used in an effort to tether the assembly scaffolds into chromosomal-level pseudomolecules. Using GBS data, a linkage map was constructed from 942 markers present in 46 scaffolds that captured approximately 50% of the sequence space. The largest 11 linkage groups constituted a 670 cM map (File S2) comprised of 898 markers among 41 scaffolds (25 of which were in the 36 N50 set and all of which were within the N85 set) that together captured 49% of the sequence space. In every case, markers within a given scaffold mapped to a single linkage group. However, within LGs 1, 3 and 5, markers were ordered on the genetic map such that sections of different scaffolds interleaved with other scaffolds (Table S2). Comparative mapping ordered 3,839 scaffolds which were anchored across all 11 *P*. *vulgaris* chromosomes (File S3). Only 16 of these scaffolds were present on nine of the *P*. *lunatus* linkage groups, which constituted a separate fraction of the assembly (~230 Mb or 37% of the assembled genome). Scaffolds anchored to two of the *P*. *vulgaris* chromosomes mapped to different linkage groups. Therefore, this study could not resolve ordering discrepancies as they could arise from errors in the genetic map, comparative map or the assembly. Additional studies will be required to construct pseudomolecules for the Bridgeton genome. Nevertheless, next we describe how combining all of the map data helped to identify a section of the genome associated with variation in partial and complete disease resistance phenotypes.

We mapped a QTL on LG 9 (*PlPp_LG9.1*) that explained ~50% of the phenotypic variation in partial resistance to *P*. *phaseoli* (a slow downy mildew lesion-development phenotype). Previously, using a different population, we used bulked segregant analysis (BSA) to map a race-specific major effect gene also associated with resistance to *P*. *phaseoli* (Mhora et al. 2016). Anchoring marker sequences from QTL and BSA mapping onto the physical map showed that both of these resistance loci reside at the same region of the genome indicating these resistance loci are linked or that *PlPp_LG9.1* is a weak allele of the race-specific resistance gene.

Markers present in scaffold 25456 were associated by both QTL and BSA mapping. Two additional scaffolds (3610 and 974) with markers detected by BSA were absent from the linkage map used for QTL analysis (due to marker QC filtering [scaffold 3610] and failure to join a linkage group [scaffold 974]), yet scaffold 974 contained markers detected by Mhora et al. (2016) showing the strongest association. Comparative mapping was used to delimit the physical section containing both resistance loci, which required a relatively low sensitivity threshold for chromosomer (-r 1.01; default is 1.2) to incorporate the three scaffolds (File S4). An additional 12 scaffolds also mapped to the region. Based on these results, genetically-associated markers spanned ~1.3 Mb across 15 scaffolds, flanked by scaffolds 3610 and 25456. Consistent with previous findings (Mhora et al. 2016), the region was homologous to a 2.5 Mb section on the short arm of chromosome 4 of *P*. *vulgaris*.

### Genome-wide annotation

Augustus predicted 64,541 genes from the masked assembly (File S5). As is typical of draft genome assemblies, this is likely an over-estimation of the true gene number in *P. lunatus*. Inflated gene counts can result from assembly fragmentation (a single gene sequence spread across multiple contigs) and failure to join distant exons together in a single transcript (Denton *et al*. 2014). The BUSCO analysis also suggests over-estimation of the true gene number: approximately 13% of the core genes were either duplicated or fragmented.

We compared our results to the genome of *P. vulgaris*, a close relative of *P. lunatus* (Delgado-Salinas et al. 2006; Bitocchi et al. 2017). The *P. vulgaris* genome has an estimated size of 587 Mb, with 98% of the sequence anchored on 11 pseudomolecules (Schmutz *et al*. 2014). After extensive annotation, Schmutz et al. (2014) predicted a final gene set consisting of 27,197 protein-coding genes with 31,638 protein-coding transcripts. We found that ~50% (32,505) of the *P. lunatus* proteins had hits to *P. vulgaris*, which encompassed 96% of the *P. vulgaris* proteins (File S6). Of the 32,036 predicted *P. lunatus* proteins without a match in *P. vulgaris*, 1,356 had hits to other Viridiplantae species. A further 2,363 had hits to non-Viridiplantae entries in the nr database, and these were predominantly to bacteria. There were 903 hits to Enterobacteriaceae, which are the dominant members of many phyllosphere bacterial communities (Cernava *et al*. 2019), 359 hits to Rhizobiaceae, a family of plant-associated bacteria (Spaink *et al*. 2012), and 254 hits to Flavobacteria, which are thought to contribute to plant growth and protection (Kolton *et al*. 2016). Only 132 of the non-Viridiplantae nr hits were to eukaryotes (90 metazoa, 31 fungi, 6 Alveolata, 2 Stramenopiles, 2 Euglenozoa, 1 Pyrenomonadales). The remaining 28,317 proteins had no hits to any sequence in the nr database. On average, the no-hit proteins were shorter (mean of 237 aa) than those with hits (mean of 514 aa).

### Resistance genes

One of our objectives in sequencing the lima bean genome was to catalogue resistance genes (R genes) commonly associated with race-specific disease resistance (citation). The majority of R genes are defined by two major domains: a centrally located nucleotide binding site domain, which has ATPase activity (NB-ARC) and a C-terminus leucine rich repeat domain (NB-LRR) (van Ooijen et al. 2008; Takken and Goverse 2012). The *P. vulgaris* genome contains 376 predicted NB-LRR genes, while *P. lunatus* Bridgeton assembly contained 206. As in many other species, NB-LRR genes tend to be clustered. The *P. vulgaris* genome has three particularly large clusters containing more than 40 NB-LRRs at the ends of chromosomes 4, 10 and 11 (Schmutz et al. 2014 [Supplementary Figure S16]). The segment on chromosome 4 is homologous with the locus described in this study that associated with multiple forms of downy mildew resistance. Out of the 15 scaffolds mapped to this region, the two scaffolds with the strongest associations (scaffolds 974 and 2456) were the only ones with (NBS-LRR) genes (File S7). However, this included only nine of the total R-genes across scaffolds. As noted above, comparative analysis with flanking marker sequences suggested that 50% of the associated region is anchored or captured by the lima bean assembly. Thus, the current draft assembly is useful for genome-wide characterization, but additional sequencing and chromosome anchoring work are required to further dissect some specific genomic regions.

## CONCLUSIONS

Closing a major gap in resources for lima bean, this study reports a near-complete reference genome for *P*. *lunatus* (Table 1). De novo protein predictions showing high similarity to 96% of the encoded genes for *P*. *vulgaris* (common bean) helped to identify 32,505 de facto genes of lima bean. The sequenced variety, Bridgeton, is the primary founder of modern cultivars for the Mid-Atlantic. In this region of the USA diseases limit the production of lima bean. Using the genome assembly, we consolidated genetic map data for loci associated with partial, race non-specific resistance (T. Evans, pers. comm.) and complete, race-F specific resistance to *Phytophthora phaseoli*, the causal agent of downy mildew. These loci co-localize in a segment that aligns to a section of chromosome 4 in common bean which is enriched with canonical R-genes and associations with resistance to different diseases – the B4 R-gene cluster (David et al. 2008; David et al. 2009). The full content of R-genes in this region for lima bean remains unresolved. However, inheritance studies in lima bean also support linkage between race-F and race-E specific resistance genes to *P. phaseoli* (Santamaria et al. 2018). Taken together, the reference genome of lima bean enabled identification of a putative ‘hotspot’ for the evolution of resistance alleles in lima bean. Improved scaffolding and more complete genome coverage is needed to genetically dissect this region, as well as to further examine evolutionary events inferred to underlie formation of the B4 R-gene cluster (David et al. 2009).

## ACKNOWLEDGEMENTS

The authors gratefully acknowledge funds from the Delaware Department of Agriculture, grant award number 58-8042-7-079 to NM Donofrio, TT Mhora and TA Evans, and 15-SCBGP-DE-0028 to EG Ernest and RJ Wisser. Some of this work was conducted on the BIOMIX compute cluster, made possible through funding from Delaware INBRE (NIGMS P20GM103446), the State of Delaware, and the Delaware Biotechnology Institute.

---

**Table S1.** Raw Sequencing Data

| Library type | Reads Length | Insert Size | Number of libraries | Genomic Coverage |
| --- | --- | --- | --- | --- |
| PCR-free | 2x260bp | 470 | 2 | X133 |
| PCR-free | 2x160bp | 800 | 2 | X36 |
| Mate-Pair | 2x160bp | 3000 | 2 | X67 |
| Mate-Pair | 2x160bp | 6000 | 2 | X58 |
| Mate-Pair | 2x160bp | 9000 | 2 | X57 |

**Table S2.** Scaffolds in the genetic map with markers differing in order on the physical map.

| Linkage Group | Interleaving Scaffolds | Max Distance (cM)<sup>1</sup> | Scaffold<sup>2</sup> |
| --- | --- | --- | --- |
| 1 | s719; s5841; s18236; s3927 | 33.19 | s18236 |
| 3 | s10495; s10613; s10634; s12141;<br>s15450; s22322; s26009 | 6.09 | s15450 |
| 5 | s9349; s7806; s13072; s10851 | 9.21 | s13072 |

<sup>1</sup>The maximum genetic distance between markers on the same scaffold<sup>2</sup> separated by other scaffolds.

